# Elevated expression of LGR5 and WNT signalling factors in neuroblastoma cells with acquired drug resistance

**DOI:** 10.1101/449785

**Authors:** Svetlana Myssina, John Clark-Corrigall, Martin Michaelis, Jindrich Cinatl, Shafiq U. Ahmed, Jane Carr-Wilkinson

**Affiliations:** Faculty of Health Sciences and Wellbeing, University of Sunderland, United Kingdom; School of Biosciences and Industrial Biotechnology Centre, University of Kent, Canterbury, United Kingdom; Institut für Medizinische Virologie, Klinikum der Goethe-Universität, Frankfurt am Main, Germany

**Keywords:** LGR5, LRP6, WNT pathway, neuroblastoma, acquired drug resistance

## Abstract

Neuroblastoma (NB) is the most common paediatric solid cancer with high fatality, relapses and acquired resistance to drug therapy. The clinical challenge NB poses requires new therapeutic approaches to improve survival rates.

The WNT signalling pathway is crucial in embryonic development but has also been reported to be dysregulated in glioblastoma, ovarian, breast and colorectal cancer. LGR5 is a receptor which potentiates the WNT/β-catenin signalling pathway, hence contributing to cancer stem cell proliferation and self-renewal. LGR5 has been reported to promote both development and survival of colorectal cancer and glioblastomas.

Our previous study illustrated that LGR5 is associated with aggressiveness in NB cell lines established at different stages of treatment. Following these findings, we investigated whether LGR5 is involved in acquired drug resistance via the WNT pathway in NB cell lines.

Cell lines in this study have an acquired drug resistance to vincristine (VCR) or doxorubicin (DOX).

In this study, we showed LGR5-LRP6 cooperation with enhanced expression of both proteins in SHSY5YrVCR, IMR32rDOX, IMR5rVCR and IMR5rDOX NB cell lines compared to paired parental cells. We also found elevated expression of β-catenin in cell lines with acquired drug resistance is indicative of β-catenin-dependent WNT signalling.

This study warrants further investigation into the role of the WNT signalling pathway in acquired drug resistance.

## Introduction

Neuroblastoma is a paediatric malignancy that originates from multi-potent embryonic neural crest cells which give rise to the sympathetic nervous system. Development of acquired resistance following induction therapy is a problem in patients with high risk neuroblastoma (categorised by the presence of *MYCN* amplification or patients with metastatic disease over 18 months), with around 50% of children relapsing with disease that is chemoresistant (Amoroso, 2018).

The cancer stem cell hypothesis, originally proposed by (Hamburger, 1977), postulates that a sub-population of cancer cells exhibit stem-like properties i.e. self-renewal and differentiation potential (Dick, 2008; Shackleton, 2009). Conventional cytotoxic agents, used in the treatment of cancers including childhood cancer, target cells which are rapidly dividing, however it has been shown the cancer stem cells are resistant to those therapies. In the last decade emerging evidence has identified the presence of cancer stem cells in tumours from several adult and paediatric cancers, including neuroblastoma (Gasch, 2017).

A study by Hansford (2007) showed the presence of tumour initiating cells, a sub-population of cells, which exhibit stem cell properties that possess the ability to initiate tumour growth (Hansford, 2007). Many stem cell markers are emerging as prognostic factors for carcinogenesis, tumour aggressiveness and therapeutic resistance. Singh’s study was one of the first to describe the presence of cancer stem cells in solid brain tumours showing that CD133+ cells possessed the capacity for tumour initiation, self-renewal and potential tumour hierarchy; in which CD133+ may generate CD133-cells (Singh, 2004). A study by Sartelet (2012), determined CD133-tumours had significantly better 3-year event survival than their CD133+ counterparts. They also found that treatment of unsorted NB cells with three anticancer drugs *in vitro*, significantly enriched the CD133+ subpopulation. Whilst CD133 and others have been described as stem cell markers associated with poor prognosis and heightened chances of therapeutic resistance, the markers do not work alone. A recent study in Hebei, China supported findings by Singh (2004), which investigated tumour cells from 50 patients at different stages of NB. These cells which were CD133+ could form differentiated neurospheres from a single tumour cell. Their study also illustrated that patients with a CD133+ and gross MYCN amplification had significantly poorer prognoses than patients with CD133-/low-MYCN amplification (Zhong, 2018). Furthermore Rosiq et al (2018), showed that CD133 and LGR5 expression were useful prognostic factors in colorectal cancer, they determined significantly increased expression of both CD133 and LGR5 in late stage carcinoma compared to normal colon cells (Rosiq, 2018).

The leucine-rich repeat-containing G-protein-coupled receptor 5 (LGR5) was originally reported as a stem cell marker in cells of the small intestine and colon (Barker, 2007) and later in mammary glands (Kumar, 2014). Recently, elevated LGR5expression has been observed in stem cells of the stomach (Leushacke, 2017) and kidney (Cao, 2017). Functionally, LGR5 modulates WNT signalling in the presence of the ligand R-spondin (RSPO) (Kumar, 2014). Recent research has shown that LGR5 is expressed in cancer stem cells (CSCs) and when overexpressed LGR5 can potentiate the effect of WNT/β-catenin which are key drivers of oncogenesis (Schuijers and Clevers, 2012; de Lau, 2014).

Other studies in papillary thyroid cancer have shown that elevated expression of LGR5 is associated with tumour aggressiveness (Michelotti, 2015). Elevated LGR5 expression is also linked to cancer stem cells in childhood cancers such as paediatric acute lymphoblastic leukaemia (ALL) (Cosgun, 2017) and Ewing’s Sarcoma (Nakata, 2013; Scannell, 2013). Furthermore recent studies have found elevated LGR5 expression levels in several other types of adult cancers including colorectal (de Sousa e Melo, 2017), cervical (Chen, 2014) and ovarian (McClanahan, 2006).

LGR5 is emerging in literature with association to a myriad of different complications in cancer; it has been identified as a marker of malignancy in colorectal cancer (van de Wetering, 2002; Barker, 2007). More recently LGR5 has been shown to be associated with chemoresistance in cervical cancer (Cao, 2017), and participates in carcinogenesis and stemness maintenance in breast cancer (Yang, 2015).

In light of these studies, and our previous study implicating LGR5 and tumour aggressiveness, it is conceivable that there is a link between elevated LGR5 expression and the development of an aggressive phenotype including acquisition of drug resistance in tumour cells.

Doxorubicin and vincristine are cytotoxic drugs used in the treatment of neuroblastoma of high risk at diagnosis (Park, 2008; Wagner, 2009). However, treatment with these drugs in patients with high risk neuroblastoma often leads to the development of acquired drug resistance and subsequently enhanced likelihood of relapse with aggressive disease (Kotchetkov, 2003). Acquired drug resistance is a major obstacle to treatment and accompanies relapsed disease, resulting in poor survival rates. This, gaining, insight into the mechanisms underlying acquired drug resistance in neuroblastoma is important in order to develop more effective therapies for this disease

Our previous study showed increased expression of LGR5 in aggressive relapsed neuroblastoma cell lines compared with non-aggressive cell lines (Forgham, 2015). Here we investigated the role of the WNT signalling pathway, including up and down stream regulators in paired NB cell lines with acquired resistance to therapy. We investigated both protein and mRNA expression of LGR5 and associated up- and downstream WNT pathway regulators, in paired parental versus drug-resistant NB cell lines, to determine their role in the development of acquired drug resistance.

## Methods

### Cell culture

The drug-adapted cancer cell lines SHSY5Y, IMR5, IMR32, NGP were derived from the Resistant Cancer Cell Line (RCCL) collection (www.kent.ac.uk/stms/cmp/RCCL/RCCLabout.html). Cell lines were derived to acquire drug resistance using 10ng of Vincristine SHSY5YrVCR, IMR5rVCR, IMR32rVCR, NGPrVCR or 20ng of Doxorubicin SHSY5YrDOX, IMR5rDOX, IMR32rDOX, NGPrDOX. Cell lines were grown in Iscove’s Modified Dulbecco’s Medium (Sigma, UK) supplemented with 10% FCS, primocin 100 ug/ml (InvivoGen, UK) and 4mM Glutamine at 5% CO_2_ and 37°C. IMR32rDOX cells were seeded on Corning^®^ Matrigel^®^ Basement Membrane Matrix (BD, UK). Cells were grown at 80% confluence and passaged regularly.

### Western blotting

Cells were lysed with PhosphoSafe Extraction Reagent (Millipore, UK) and total protein stored at −80°C. Standard western blotting procedure was performed using Bio-Rad equipment. 20ug of total protein were loaded per well and membrane was probed with LGR5 (21833-1-AP, Proteintech, UK), phospho-LRP6 (phospho S1490, Abcam, UK), β-catenin (H-102) (Santa Cruz Biotechnology, sc-7199), c-Jun (N) (Santa Cruz Biotechnology, sc-45) and GSK-3α/β (0011-A) (Santa Cruz Biotechnology, sc-7291) antibody. Antibodies were diluted 1:1000 in 5% BSA solution. GAPDH used as a housekeeping control.

### Quantitative Real Time PCR

Total RNA was purified with QIAGEN RNA extraction kit (QIAGEN, UK) and first strand DNA synthesised with iScript^™^ cDNA synthesis kit (Bio-Rad, UK). Expression of LGR5 mRNA was estimated using TaqMan R gene expression assay primers and probes LGR5 (Hs00173664_ml); GAPDH (Hs0278991_g1); DKK1 (Hs00183740_m1) (Life Technologies, UK). Quantification was performed on the Applied Biosystems^®^ ViiA^™^ 7 Real-Time PCR System (Applied Biosystems, UK). Gene expression was calculated relative to GAPDH housekeeping gene using relative delta delta CT.

### Statistical analysis

Paired t-tests were performed using SPSS software and statistically significant differences accepted if p < 0.05.

## Results

### Elevated expression of LGR5 and β-Catenin in a sub-set of neuroblastoma cell lines with acquired resistance

Quantitative Real Time PCR (qRT-PCR) was performed to determine LGR5 mRNA expression levels in our panel of 4 paired parental and resistant neuroblastoma cell lines, resistant to either vincristine or doxorubicin. Elevated LGR5 mRNA expression was observed in SHSY5YrVCR (p = 0.02) and IMR32rVCR (although not significant, p = 0.13) compared with the corresponding SHSY5Y and IMR32 parental cells. Conversely, both NGP resistant cell lines and SHSY5YrDOX, show downregulated LGR5 gene expression (Figure 1). Interestingly in the MDM2 amplified NGP cell line we observed no changes in LGR5 expression in NGPrVCR (p = 0.22) and NGPrDOX (p = 0.09) compared with corresponding parental cells (Figure 1) (N = 3). β-catenin, a down-stream regulator of the WNT signalling pathway, showed elevated expression in both SHSY5YrVCR and SHSY5YrDOX compared with parental SHSY5Y cells. In contrast β-catenin expression was reduced in NGPrVCR and NGPrDOX compared with parental NGP cells (Figure 2) (N = 3).

**Figure 1.**
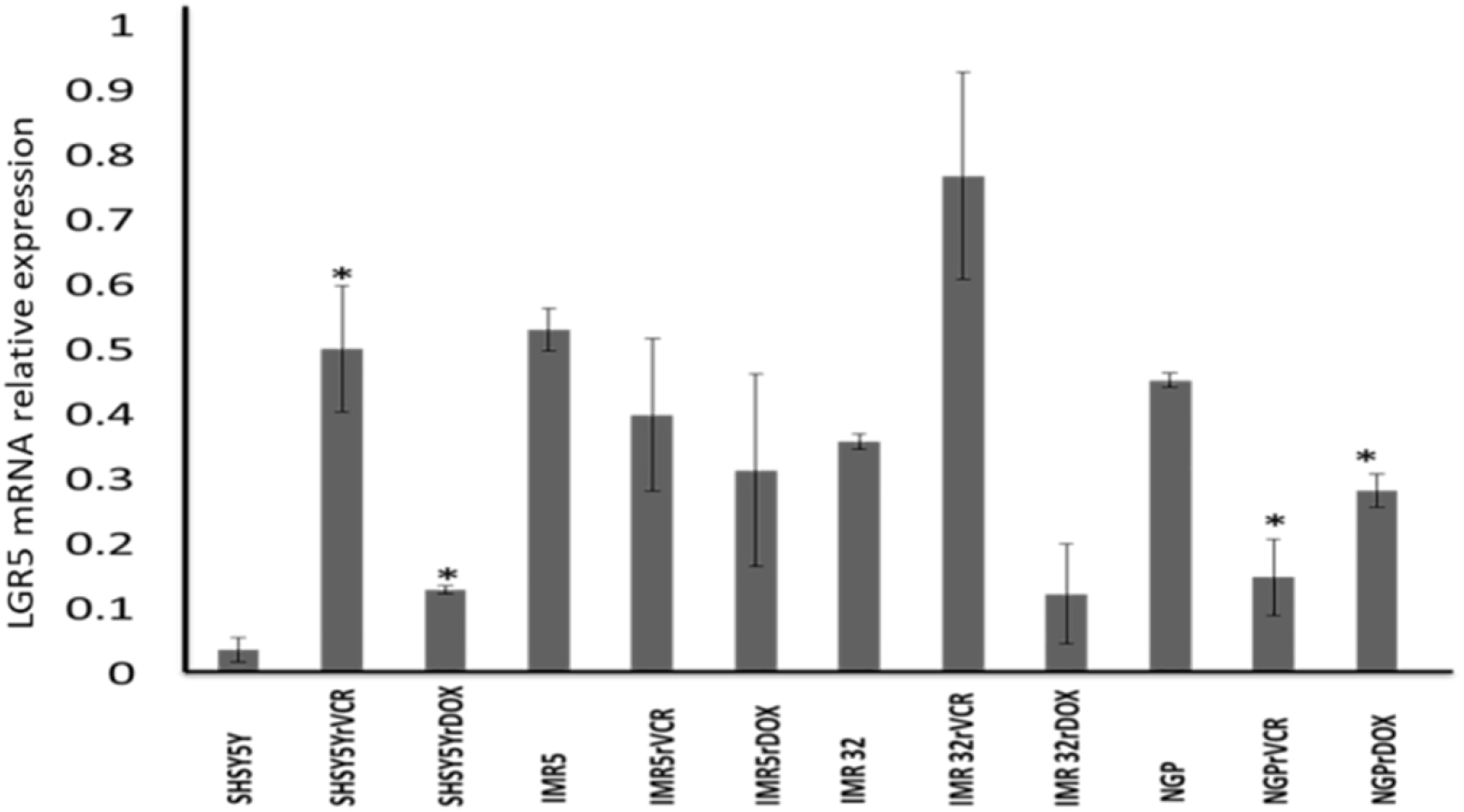
Real time Q-PCR showing LGR5 mRNA expression in neuroblastoma parental and drug-resistant cell lines. Q-PCR analysis of gene expression of paired SHSY5Y, IMR5, IMR32 and NGP cell lines (N=3).

**Figure 2.**
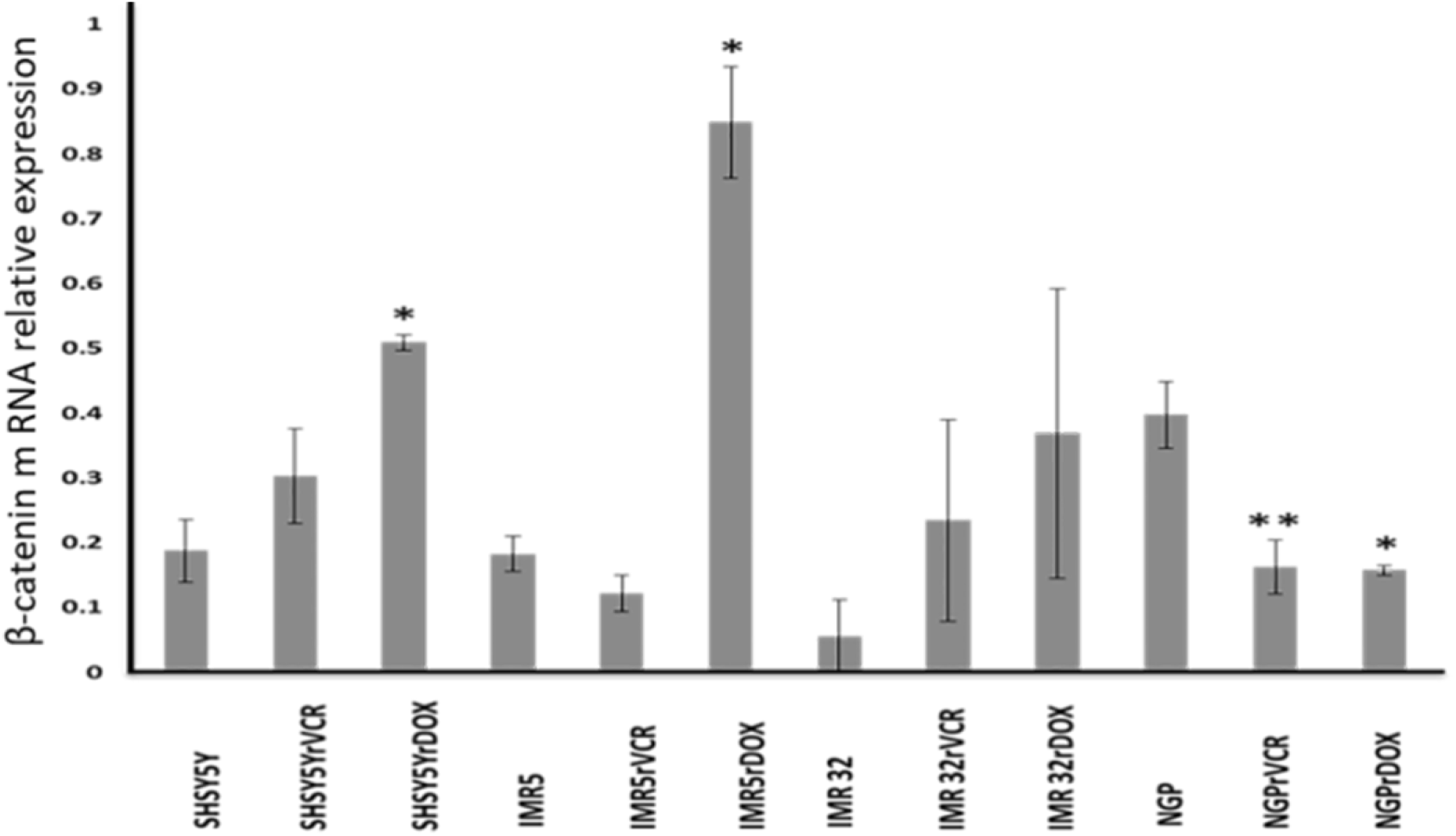
Real time Q-PCR results showing mRNA expression of β-catenin gene relative to GAPDH (P < 0.05*, P < 0.01**).

### LGR5 Protein expression in paired parental and drug resistant cell lines

To further explore the link between aggressiveness of neuroblastoma and elevated LGR5 expression (Forgham, 2015), we investigated protein expression of upstream and downstream proteins in the WNT/β-catenin signalling pathway, in addition to LGR5 in four paired parental and drug resistant cell lines. Elevated LGR5 expression was reported in 4 paired cell lines; SHSY5YrVCR (P <0.01 * *), IMR5rVCR and IMR5rDOX (P <0.001 * * *), as well as in IMR32rVCR (P <0.05 *) cell lines (Figure 3B). Elevated expression was also observed in the SHSY5YrDOX cell line however, this was not significantly different from parental cells. Conversely, IMR32rDOX showed significantly decreased LGR5 protein expression, compared with parental cells (Figure 3).

**Figure 3.**
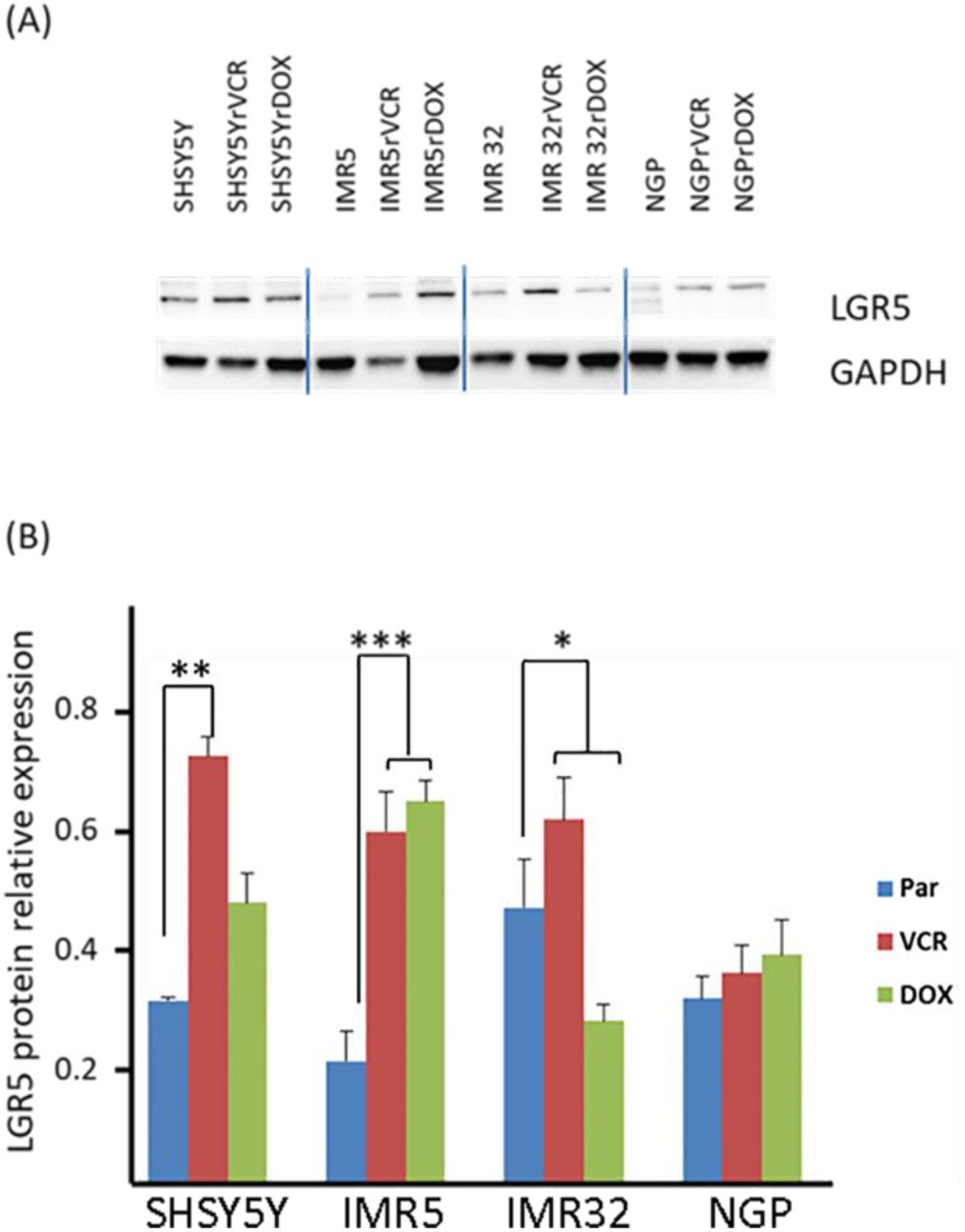
(A) LGR5 protein expression in paired parental drug resistant neuroblastoma (NB) cell lines. Western blotting was performed on SHSY5Y, IMR5, IMR32 and NGP NB cell lines using LGR5 antibody. GAPDH is used as housekeeping protein control. (B) Relative expression of LGR5 protein. Par – parental cell line; VCR – cell lines resistant to 10ng vincristine; DOX – cell lines resistant to 20ng doxorubicin (n=4). P < 0.05*, P < 0.01**, P < 0.001***.

### Upstream and downstream WNT signalling regulator analysis

Subsequently, analysis of WNT signalling upstream and downstream of LGR5 was conducted, focussing on LRP6 and GSK3β. Phosphorylation of LRP6 is indicative of WNT pathway activation (Yao, 2017). Significantly higher levels of pLRP6 expression was correlated with elevated LGR5 in all 4 cell lines; SHSY5YrVCR, IMR5rVCR and IMR5rDOX, IMR32rVCR (P < 0.01**, paired t-test).

Significantly elevated LGR5 protein expression was shown in IMR5 alongside increased protein in both upstream (Figure 4A, B) and downstream (Figure 5A, B) levels in both IMR5rVCR and IMR5rDOX.

**Figure 4.**
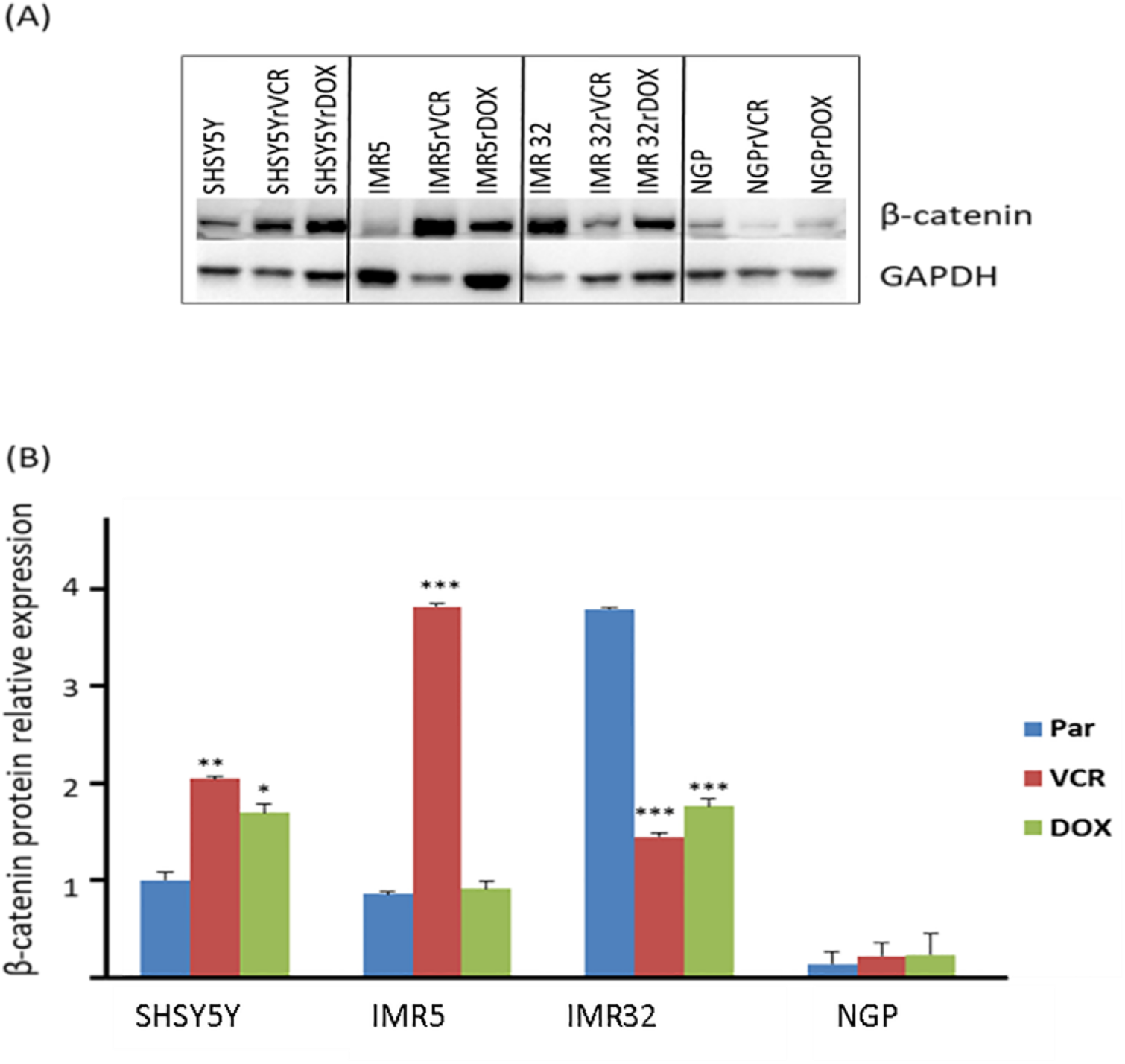
Upstream signalling of WNT pathway. (A) Activated LRP6 protein expression. Blotted membrane was incubated with anti-phospho LRP6 antibody to detect activation of LRP6 protein. (B) Relative protein expression to GAPGH. pLRP6 presented in significant higher level in SHSY5YrVCR, IMR5rVCR and IMR5rDOX, IMR32rVCR (P < 0.01**) to compare to corresponding parental cell lines. There is no significant change in NGP drug resistant cell lines (N=3).

**Figure 5.**
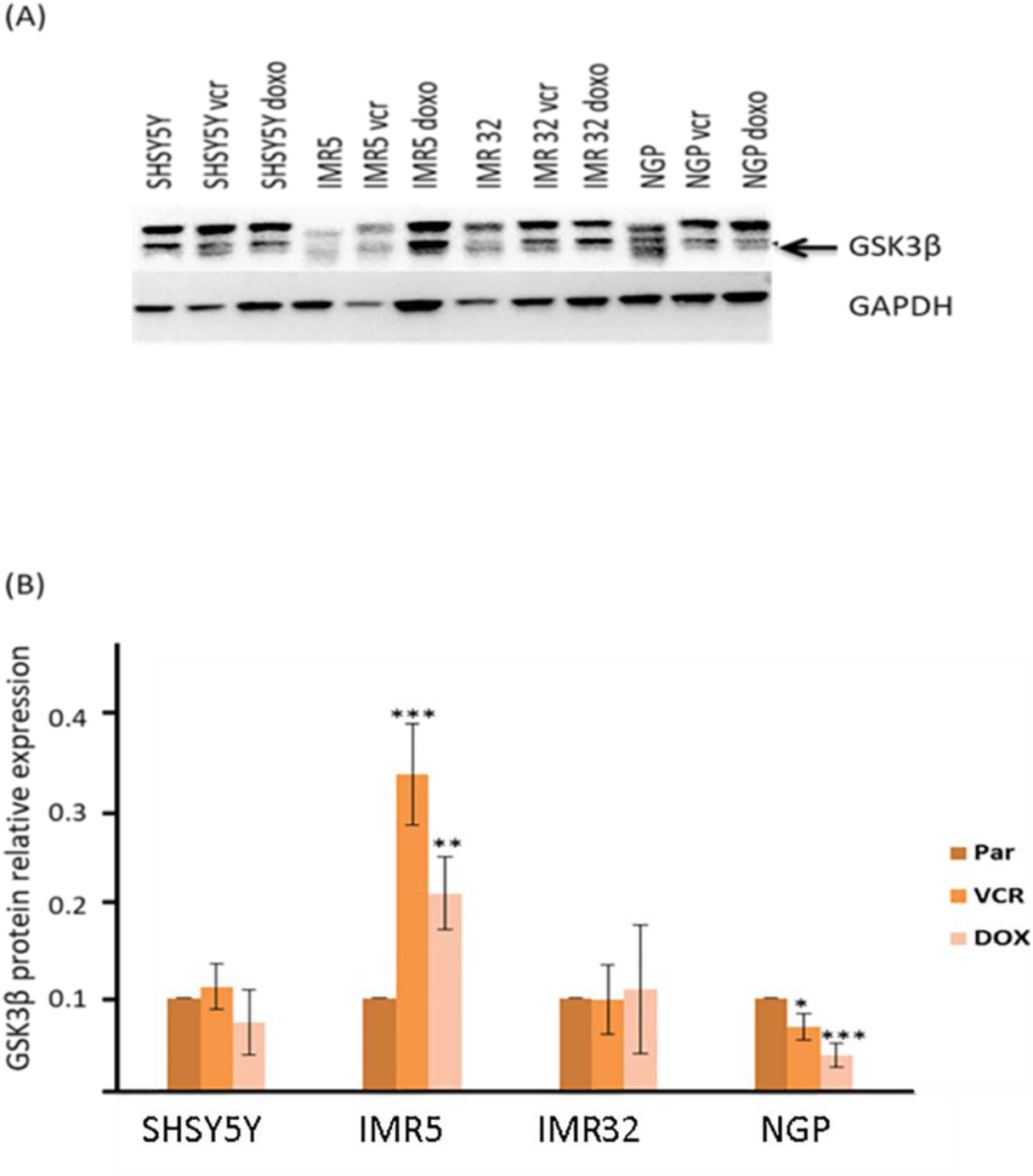
(A) Expression of GSK3 as the downstream protein of LGR5/ WNT signalling pathway in neuroblastoma cell lines with acquired drug resistance. (B) GSK3β protein expression presented as fold decrease in SHSY5Y, IMR5, IMR32 and NGP cell lines. Statistical analysis performed within cell line between parental and resistant for either VCR or DOX cell lines (P < 0.05*, P < 0.01**, P < 0.001***).

Our study observed significantly decreased levels of LGR5 expression in IMR32rDOX but no significant changes in expression to either upstream or downstream regulators. Whilst IMR32rVCR showed significant overexpression of LGR5 and LRP6, there was no significant effect on downstream expression of GSK3β (Figure 5).

Interestingly, NGPrVCR and NGPrDOX show no significant increases in expression compared to parental cells in either LGR5 or LRP6. However, there is significantly decreased expression in GSK3β in NGPrVCR (P < 0.05*) and NGPrDOX (P < 0.001***).

### β-catenin protein expression in paired parental and drug resistant cell lines

To confirm the presence of canonical WNT signalling, β-catenin protein expression was analysed across all paired cell lines. Significant overexpression of β-catenin was observed in IMR5rVCR (P < 0.001***), SHSY5YrVCR (P < 0.01**) and SHSY5YrDOX (P < 0.05*). Conversely, IMR32rVCR and IMR32rDOX both showed significant down regulation in β-catenin (P < 0.001***) (Figure 6).

**Figure 6.**
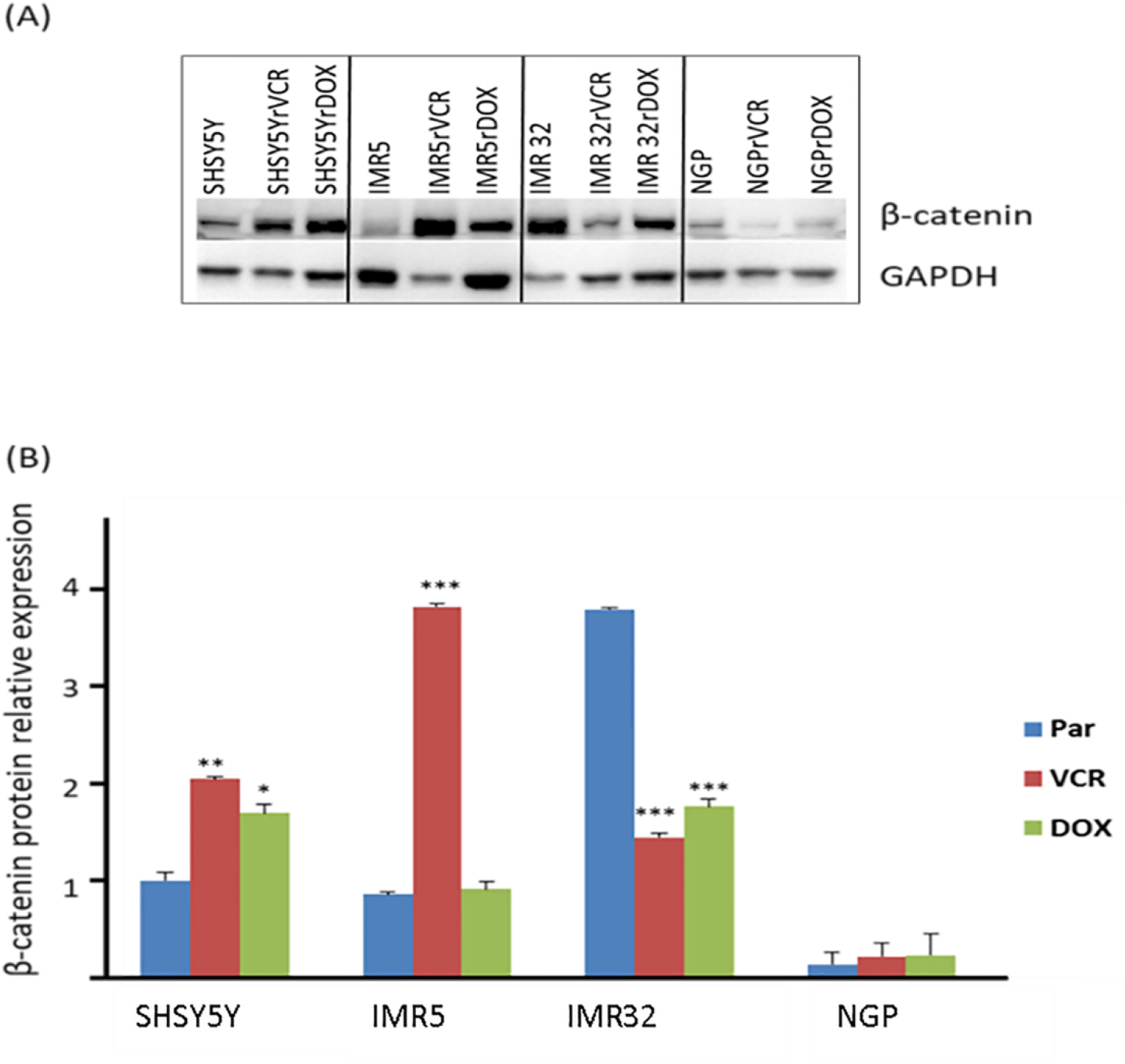
β-catenin protein expression. (A) Western blotting has shown changes in expression which quantitatively presented in (B) as relative increase in β-catenin protein expression (N=3) in drug resistant neuroblastoma cell lines SHSY5Y, IMR5, IMR32 and NGP; VCR – cell lines resistant to vincristine, DOX – cell lines resistant to doxorubicin (P < 0.05*, P < 0.01**, P < 0.001***).

## Discussion

There is a growing body of evidence which reports increased expression of LGR5 in several cancers, and its involvement in therapeutic resistance. In this study we investigated the expression of LGR5 in cell lines with acquired drug resistance. Furthermore, we also investigated genes involved in the WNT/β-catenin pathway and evaluated upstream and downstream signalling of LGR5.

Our previous study (Forgham, 2015) reported high LGR5 protein expression in parental SHSY5Y, findings which also supported observations made by Vieria (2015). Further elevation of LGR5 protein levels in vincristine-resistant SHSY5Y cells highlight a potential role for LGR5 in acquired drug resistance.

Although IMR32rVCR have shown a corresponding increase in both LGR5 and LRP6 protein expression, similar changes were not observed in IMR32rDOX. In fact, generally, LGR5 and its upstream and downstream WNT pathway proteins showed significantly higher expression in VCR-resistant cells than DOX-resistant cells (Figure 3-6).

The mechanisms in which the drugs act explain the contrasting behaviour seen in DOX- and VCR-resistant cells. Doxorubicin caused early activation of p53 in cancer cells that leads to caspase-3 dependent apoptosis (Wang, 2004).

However, vincristine acts by perturbation of mitosis via inhibition of microtubule formation in spindle and arrests cell cycle in metaphase preventing cell division. Aggressive childhood neuroblastoma has been proved to lack the important mechanism of apoptosis, caspase-8 (Teitz, 2001).

Neuroblastoma cell lines are heterogeneous containing neuronal N-type cells and adherent mesenchymal cells (Walton, 2004), and therefore express different characteristics (Corey, 2010). Hopkins-Donaldson (2002) reported that the IMR32 cell line is a caspase-8 silenced N-type cell line and that dox-induced death in N-type cells was caspase-independent. In addition they stated that DOX-induced death in S-type cells gave rise to apoptotic nuclei, whereas, in N-type cells nuclei were non-apoptotic (Hopkins-Donaldson, 2002). Simultaneously, we expected the same pattern in LGR5 and pLRP6 expression in all cell lines as they all are N-type.

Versteeg’s group (Van Groningen, 2017) have shown that most neuroblastoma cell lines include undifferentiated mesenchymal cells or committed adrenergic cells which can interconvert and resemble cells from different lineage differentiation stages. According to the study, mesenchymal cells are most chemoresistant (Van Groningen, 2017). Cell lines used in this study containing two different populations of cells however, percentage of mesenchymal cells may vary in different cell lines, thus resulting in different chemoresistant mechanisms. Also, it was reported that IMR5 and SHSY5Y lack oligonucleosomal DNA fragmentation during apoptosis (Yuste, 2001). Therefore, because these two cell lines displayed similar behaviour in apoptosis, we expected similar changes in LGR5 expression. Surprisingly, we observed conflicting results; as IMR32 and SHSY5Y DOX-resistant cells presented low protein expression, IMR5rDOX had in fact increased levels of LGR5 and pLRP6.

We observed a strong correlation between elevated LGR5 expression and increased phosphorylation of LRP6. Thus, suggesting that there is a cooperative relationship between LGR5 and pLRP6, resulting in WNT pathway activation in cell lines with acquired drug resistance. LRP6 has been reported as an activation molecule for WNT pathway and shown to play a role in carcinogenesis (Lu, 2011; Arensman, 2015; Liang, 2011). Considering the findings observed in our study there is strong evidence of the involvement of the WNT signalling pathway activation in acquired drug resistance.

LGR5 has been associated with a number of signalling pathways in several cancers; both adult and paediatric, resulting in increased tumour aggressiveness. Recently, LGR5 has been reported to promote cell-cell adhesion in stem cells and colon cancer cells via the IQGAP1-Rac1 pathway (Carmon, 2017). A study by Lawlor’s group showed that LGR5 potentiates WNT/β-catenin signalling in Ewing sarcoma (Scanell, 2013). A neuroblastoma study (Vieira, 2015) proposed that LGR5 was involved in MEK/ERK signalling.

In light of those studies, our findings suggest that in drug resistant NB cells; LGR5 potentially acts through a number of different signalling pathways, as not all of the drug resistant cell lines studied have elevated downstream WNT signalling protein expression. Peng et al (2014) has reported that proliferation and differentiation of mesenchymal stem cells are regulated by WNT/β-catenin signalling. Our results proposed that canonical WNT signalling is more likely to be involved in acquired VCR-resistance as there appears to be a correlation between LGR5, pLRP6 and downstream markers of WNT/β-catenin signalling.

NGP cells did not show increased expression of LGR5 (Figure 3), or pLRP6 (Figure 4) protein expression in cell lines with acquired resistance. A characteristic of the NGP cell line is MDM2 amplification, which can lead to p53 inactivation (Haupt, 1997). This MDM2-p53 interaction may inhibit LGR5 expression leading to a different mechanism of acquired drug resistance in NGP cells. Evasion of growth suppressors, particularly through mutations that cause p53 inactivation is a hallmark of cancer (Hanahan and Weinberg, 2011).

## Conclusion

Our study suggests a coordinated relationship between LGR5 and LRP6 in both parental and drug resistant neuroblastoma cell lines. Thus, we propose that the WNT signalling pathway exerts a role in the development of acquired drug resistance in neuroblastoma. Another important finding, was the differing expression levels in cell lines with resistance to different therapeutic agents. This highlights the importance of research into therapeutic resistance to several agents, as it would be naïve to assume that there is only one mechanism for acquired drug resistance. This is a novel avenue of research, hence warrants further study using primary tumour samples. Gaining insight into the mechanisms driving the development of acquired drug resistance is important in order to develop more effective therapies to improve patient survival rates and to combat the clinical challenge that is neuroblastoma.

## Acknowledgements

We would like to thank Florian Rothweiler for providing the cell lines. Research was supported by a University of Sunderland Research Beacon Grant.

